# Distribution and extent of suitable habitat for geladas (*Theropithecus gelada*) in the Anthropocene

**DOI:** 10.1101/2023.08.10.552774

**Authors:** Ahmed Seid Ahmed, Desalegn Chala, Chala Adugna Kufa, Anagaw Atickem, Afework Bekele, Jens-Christian Svenning, Dietmar Zinner

**Author notes:** Shared first authorship (Ahmed Seid Ahmed, Desalegn Chala). Correspondence Ahmed Seid Ahmed.

## Abstract

**Background:** Climate change coupled with other anthropogenic pressures may affect species distributions, often causing extinctions at different scales. This is particularly true for species occupying marginal habitats such as gelada, *Theropithecus gelada.* Our study aimed to model the impact of climate change on the distribution of suitable habitats for geladas and draw conservation implications. Our modelling was based on 285 presence locations of geladas, covering their complete current distribution. We used different techniques to generate pseudoabsence datasets, MaxEnt model complexities, and cut-off thresholds to map the potential distribution of gelada under current and future climates (2050 and 2070). We assembled maps from these techniques to produce a final composite map. We also evaluated the change in the topographic features of gelada over the past 200 years by comparing the topography in current and historical settings.

**Results:** All model runs had high performances, AUC = 0.87 – 0.96. Under the current climate, the suitable habitat predicted with high certainty was 90,891 km^2^, but it decreased remarkably under future climates, −36% by 2050 and −52% by 2070. Whereas no remarkable range shift was predicted under future climates, currently geladas are confined to higher altitudes and complex landscapes compared to historical sightings, probably qualifying geladas as refugee species.

**Conclusions:** Our findings indicated that climate change most likely results in a loss of suitable habitat for geladas, particularly south of the Rift Valley. The difference in topography between current and historical sightings is potentially associated with anthropogenic pressures that drove niche truncation to higher altitudes, undermining the climatic and topographic niche our models predicted. We recommend protecting the current habitats of geladas even when they are forecasted to become climatically unsuitable in the future, in particular for the population south of the Rift Valley.

## 1 INTRODUCTION

Climate change is triggering alterations and shifts in ecosystems worldwide, causing changes in the distribution and availability of suitable habitats for many species, including primates [1–6]. For most taxa distribution models predict substantial habitat loss by 2100 due to climate change [7]. In particular, the expected upslope shift of habitats in mountain areas is expected to lead to a reduction of suitable habitats and the extinction of range-restricted high-altitude species [8–10].

Among primates, a few species belong to such range-restricted high-altitude species, e.g., snub-nosed monkeys (*Rhinopithecus* spp.) in China and Myanmar [11, 12] and geladas (*Theropithecus gelada*) in Ethiopia [13]. Geladas are endemic to Afro-alpine grasslands of Ethiopia at elevations from 1800 m to 4400 m asl [14–17]. Three populations, whose taxonomic status is unclear, are recognized: *T. g. gelada* in northern Ethiopia, mainly in the Simien Mountains, *T. g. obscurus*, in the central highlands of Ethiopia, and a small population south of the Rift Valley in the Arsi Mountains (*T*. *g*. ssp. nov.) [17–19; Fig. 1]. Interestingly, Chiou et al. [19] found a chromosomal polymorphism in geladas that could potentially contribute to reproductive barriers between populations, which suggests specific status for the three populations (subspecies).

**Fig. 1.**
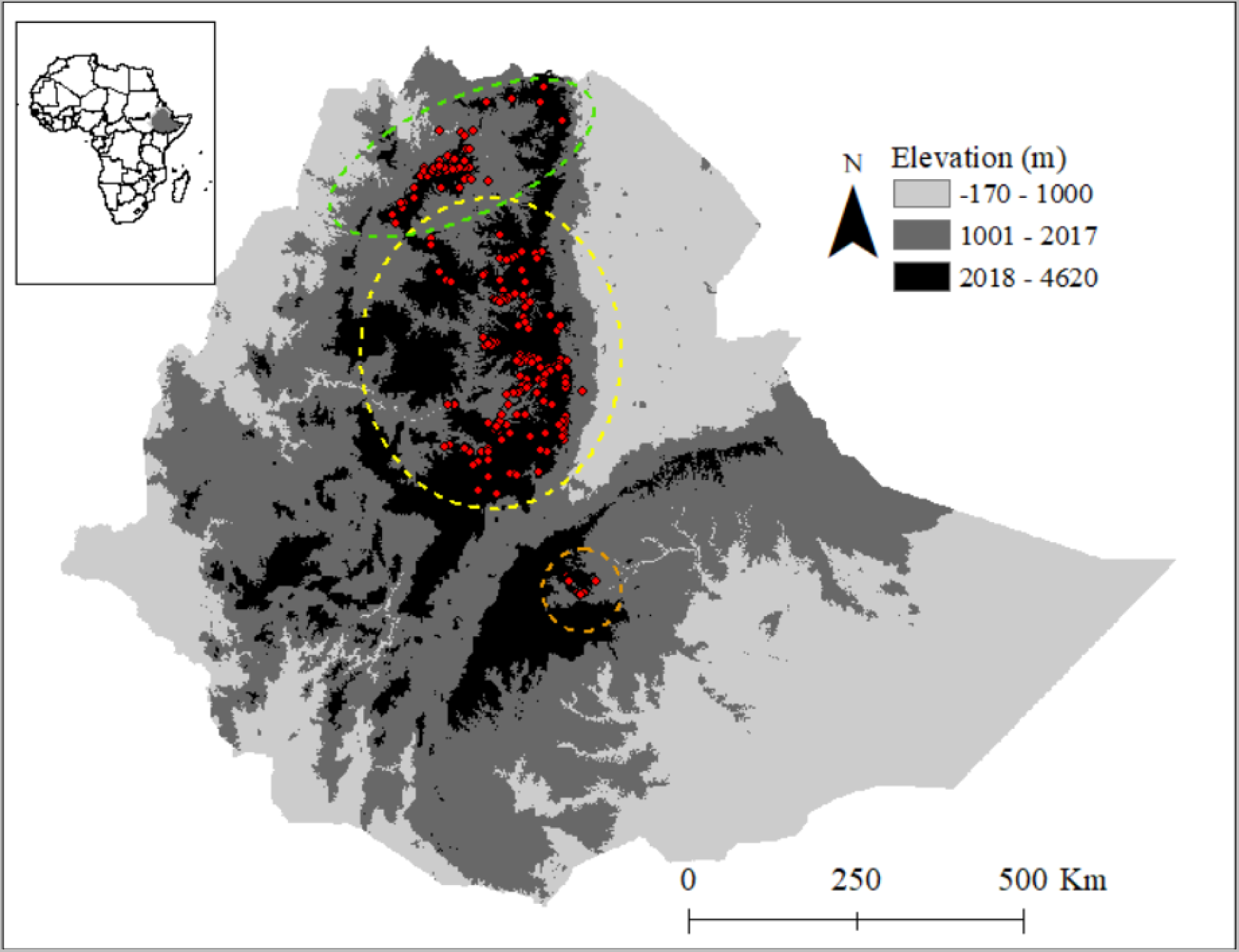
Topographic map of Ethiopia indicating the relief and geographic positions of occurrence locations of geladas (*Theropithecus gelada,* red dots) after 2000. The broken lines encircle the assumed distribution ranges of the northern (*T*. *g*. *gelada*, green), the central (*T*. *g*. *obscurus*, yellow), and the southern (*T. g.* ssp. nov., orange) populations [18, 32].

Even without taking the effects of climate change into account, the population size of geladas is generally decreasing due to the conversion of their habitat into farmland, grazing grounds for livestock, and settlements [17]. *T. g. obscurus* is listed as of Least Concern by IUCN [20], whereas *T. g. gelada* is listed as Vulnerable [21]. The conservation status of the southern population is not yet officially assessed, but it seems that this population is under particular pressure due to its already small population size and extreme anthropogenic habitat conversion [22, 23].

In general, the effects of climate change on Ethiopia’s biodiversity have not been well studied [24], but recently the effects of climate change on habitat suitability of two other endemic high-altitude species of the Ethiopian highlands have been modelled, the Walia ibex (*Capra walie*) and the giant lobelia (*Lobelia rhynchopetalum*). In both studies, significant reductions in the size of the species ranges have been projected [9, 25]. In a pioneering study on geladas, Dunbar [26] already estimated that for every 2°C rise in the mean temperature, the lowest altitude geladas may inhabit will rise by 500 m.

For adequate conservation strategies under climate change, it is essential to include information on future potential distributions of suitable habitats [24, 27]. Species distribution models (SDMs) based on current presence-absence data or presence data alone in combination with climate change models can be applied to predict the spatiotemporal changes in suitable habitats [28–31]. In our study, we applied species distribution modelling to project the distribution and extent of suitable habitats for geladas in the Ethiopian highlands under 2050 and 2070 climate change scenarios.

## 2 MATERIALS AND METHODS

### 2.1 Occurrence data

We assembled occurrence points for the three subspecies of geladas from different sources such as personal surveys (n= 396), literature [15, 32–35; see also Table S1) and from GBIF.org [36]. These occurrence data were collected after 1999 to represent the current presence data of geladas. To explore whether gelada occurrence already changed topographically (e.g., altitude of occurrence), we compared historical occurrence data [14, 33] collected before 2000 with the current data. For this comparison, we divided the historical data into data collected before 1900 and data collected between 1900 and 1999. We included only data collected after 1999 in our modelling approach. We further filtered this data by removing duplicates, and, in cases where we detected multiple occurrence points within 1 km × 1 km grid area, we included only one point. Finally, we retained 285 occurrence points for our modelling (Fig. 1). Since the number of occurrence points for each subspecies was not sufficient for proper modelling at subspecies level, we restricted our analysis to the species (genus) level.

### 2.2 Environmental variables

We initially considered 23 environmental variables for the modelling including 19 bioclimatic variables, land cover (https://cds.climate.copernicus.eu/) and vegetation type (http://landscapeportal.org/layers/geonode:veg_ethiopia), and two topographic variables (slope and slope SD). We obtained the bioclimatic variables from the WorldClim v2.1 at a spatial resolution of 30 arc seconds (~1 km^2^) [37]. Geladas frequently use more or less flat areas on plateaus for foraging and steep cliffs as a refuge from predators and as sleeping sites [16; Fig. 2]. Therefore, we added slope data. We derived the slope angle map from a digital elevation model downloaded from the Shuttle Radar Topography Mission Digital Elevation Model [SRTM DEM; 38]. In the previous study, it was shown that manually collected occurrence points for animals adapted to complex topographic landscapes tend to be confined to their foraging sites and not to places where the animals are in their inactive phases (sleeping sites) and taking refuge from predators [25]. This relationship most likely caused that slope was not found to be an important predictor variable for the Walia ibex (*Capra walie*), although this species is a steep-slope specialist [25]. Hence, topographic complexity may better predict the topographic requirements of geladas. Thus, to represent topographic complexity, we additionally computed the slope standard deviation as a proxy from pixels within a radius of three 1-km^2^ grid cells around one central grid cell for the whole landscape of the study area and used it as an additional predictor.

**Fig. 2.**
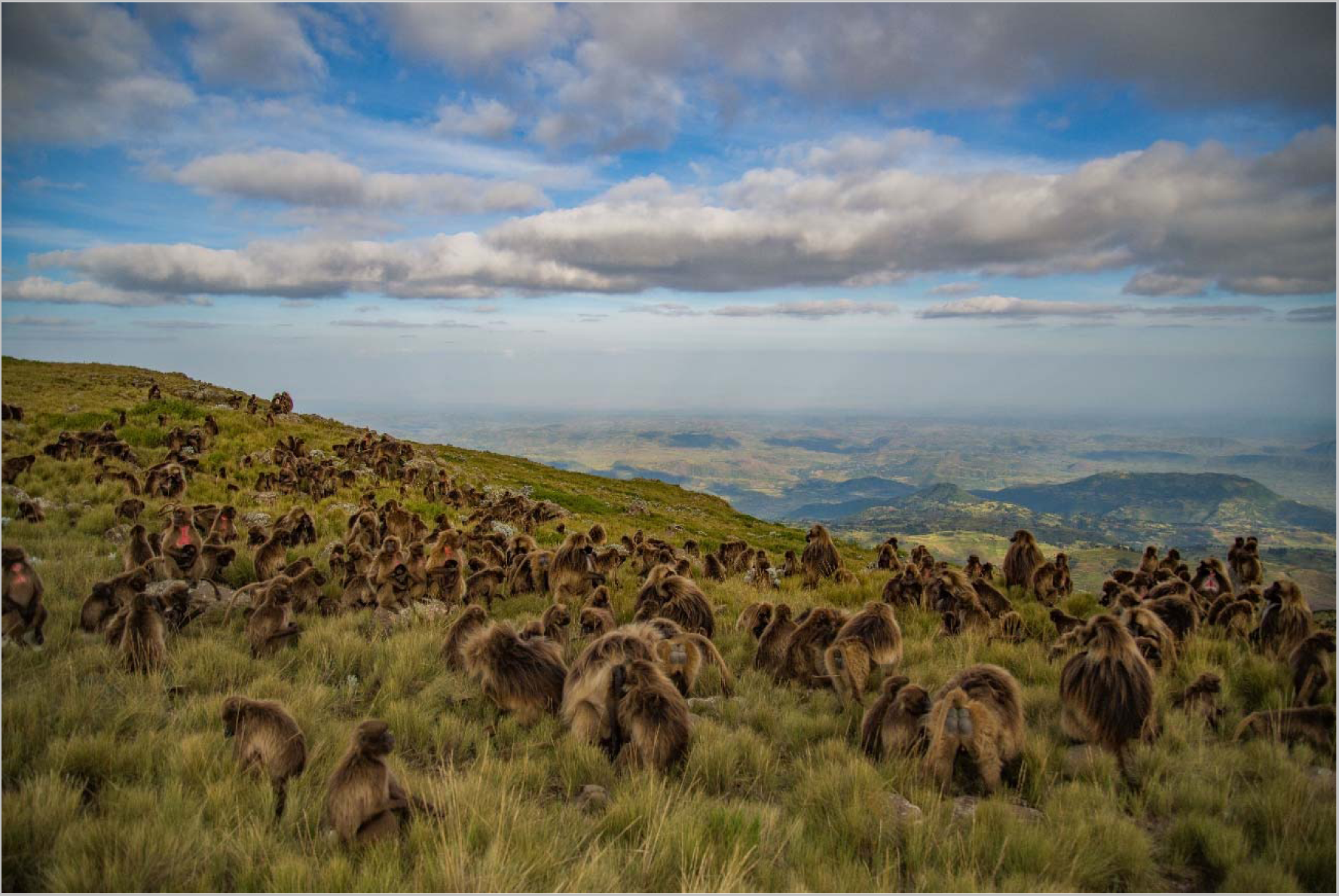
Gelada herd (*Theropithecus gelada obscurus*) in the Afro-apline grassland in the highlands of cental Ethiopia (Guassa Community Conservation Area). Photos credit - Jeffrey T. Kerby

To avoid multi-collinearity, we stacked all 23 environmental variables and extracted their values at each of the occurrence points and additionally at 10,000 randomly generated points. Based on these points, we computed Pearson’s pairwise correlations among all variables. From variables with a pairwise correlation coefficient of r > |0.8|, we retained only those variables that had the lowest variable inflation factor values, computed in the ‘USDM’ R package [39; Fig. S1]. With this procedure, we reduced the number of environmental variables from 23 to 13 for the final model run (Table 1).

**Table 1.**
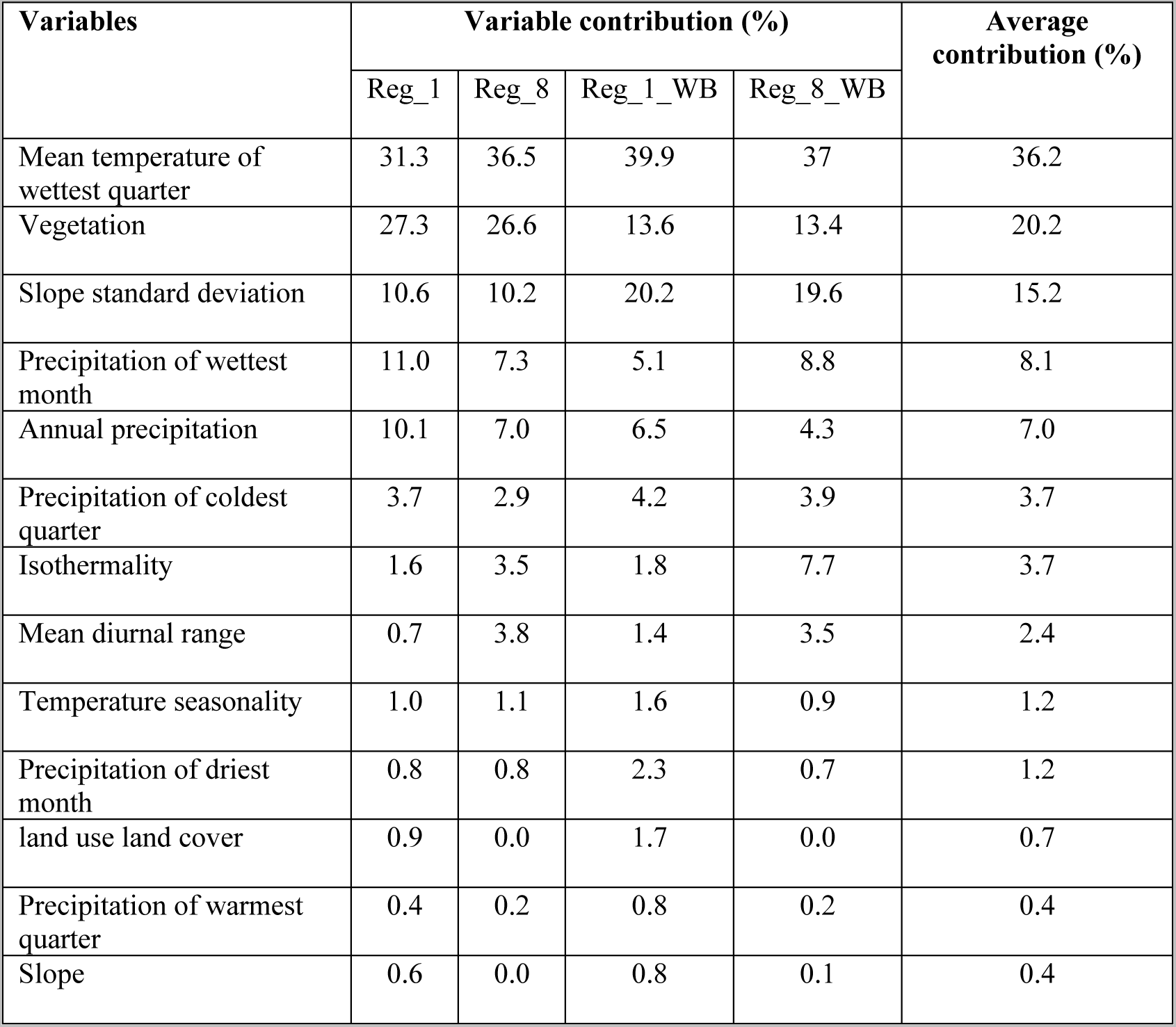
Contributions of the predictor variables to the four MaxEnt models. Reg_1: complex MaxEnt model run with pseudo-absence points generated within the elevation range of geladas; Reg_8: simple MaxEnt model run with pseudo-absence points generated within the elevation range of geladas; Reg_1_WB: complex MaxEnt model run with pseudo-absence points generated using bias file; Reg_8_WB: simple MaxEnt model run with pseudo-absence points generated using bias file.

| Variables | Variable contribution (%) |  |  |  | Average contribution (%) |
| --- | --- | --- | --- | --- | --- |
|  | Reg_1 | Reg_8 | Reg_1_WB | Reg_8_WB |  |
| Mean temperature of wettest quarter | 31.3 | 36.5 | 39.9 | 37 | 36.2 |
| Vegetation | 27.3 | 26.6 | 13.6 | 13.4 | 20.2 |
| Slope standard deviation | 10.6 | 10.2 | 20.2 | 19.6 | 15.2 |
| Precipitation of wettest month | 11.0 | 7.3 | 5.1 | 8.8 | 8.1 |
| Annual precipitation | 10.1 | 7.0 | 6.5 | 4.3 | 7.0 |
| Precipitation of coldest quarter | 3.7 | 2.9 | 4.2 | 3.9 | 3.7 |
| Isothermality | 1.6 | 3.5 | 1.8 | 7.7 | 3.7 |
| Mean diurnal range | 0.7 | 3.8 | 1.4 | 3.5 | 2.4 |
| Temperature seasonality | 1.0 | 1.1 | 1.6 | 0.9 | 1.2 |
| Precipitation of driest month | 0.8 | 0.8 | 2.3 | 0.7 | 1.2 |
| land use land cover | 0.9 | 0.0 | 1.7 | 0.0 | 0.7 |
| Precipitation of warmest quarter | 0.4 | 0.2 | 0.8 | 0.2 | 0.4 |
| Slope | 0.6 | 0.0 | 0.8 | 0.1 | 0.4 |

For the temporal projections, we used the HadGEM3-GC global circulation model (GCMs) with three shared socioeconomic pathways (SSPs): (1) the straightest emission pathway scenario (SSP 2.6), (2) the intermediate (SSP 4.5), (3) the worst (SSP 8.5) and applied them for two periods (2041–2060 [2050] and 2061–2080 [2070] [40].

### 2.3. Model fitting

We used the maximum entropy algorithm MaxEnt v3.4.4 [41] to model suitable habitats for geladas under the current climate and for the projection to future climate scenarios. MaxEnt is a commonly used algorithm to predict species distributions and is robust even with small sample sizes [29, 42, 43]. One factor that affects the model performance of MaxEnt is the spatial extent from which pseudo-absence points are taken [44–46]. Generating pseudo-absence points over larger areas that are already known to be unsuitable to the model species may exaggerate model performance. Thus, restricting the spatial extent of pseudo-absence points is important [9, 25, 47–49]. We generated 10,000 pseudoabsence points as implemented in default MaxEnt [41]. However, we restricted these points to areas where we expect suitable habitats for geladas by two approaches. First, we used a bias file [48]. The bias file works by minimizing omission (false negatives) and commission errors (false positives), which may improve the prediction performance of the model [50]. We created a bias file in ArcGIS version 10.7 by mapping species records on a 1-km^2^ grid and producing a minimum convex polygon. Second, we restricted the area to the elevation range where geladas currently are known to occur. We extracted the altitude of each gelada occurrence point and generated pseudoabsence points within 90% of the total elevation range, omitting 2.5% of lower and upper ranges.

We combined the gelada occurrence points with both datasets for pseudoabsence and used them and the values of the selected environmental variables as input into MaxEnt. We split both combined datasets into ten equal parts using a cross-validation technique and run ten replicates of two versions of the MaxEnt model [51], one simple and one complex. The complex model was run by setting the regularization multiplier value to 1 which is the default MaxEnt setting [52], and to 8 for the simple model [25]. Eventually, we run four MaxEnt models: two complex models, one using the pseudo-absence points generated using bias file and one using the pseudo-absence points generated within the elevation limits of geladas occurrence points, and two simple models using the same two datasets. For all model runs, we used 90% of the combined occurrence and pseudo absence points for model training and (10%) of the data for validation. The robustness of the models was evaluated with 5000 iterations [41, 53]. All four MaxEnt models were projected into the three emission scenarios by 2050 and 2070 (see above). We classified the output maps from all models and model projections into binary suitable/unsuitable classes using three probability threshold criteria: (1) 10 percentile logistic training threshold, which is the predicted probability at a 10% omission rate of the training data; (2) using maximum test sensitivity plus specificity, which is the probability threshold at which the sum of fractions of correctly predicted presence and pseudo-absence points is the highest; and (3) using equal test sensitivity and specificity, which is the probability thresholds at which the difference between fractions of correctly predicted presence and pseudo-absence points are the lowest. In sum, we produced 12 binary maps for the current climate (four versions of the MaxEnt model with three threshold criteria for each version; Fig. S2) and 36 binary maps for future climate (four versions of the MaxEnt model times three threshold criteria times three emission scenarios).

We ensemble the binary maps from both, current and future climate scenarios, and produced three habitat suitability classes based on agreements among the maps in predicting habitat suitability [9]: (1) highly suitable, when pixels from more than 60% of the binary maps predict habitat as suitable (≥ 8 maps for the current climate and ≥ 22 maps for future climates); (2) uncertain, when 30 – 60% of the maps predict habitat suitability (4-7 maps for current climate and 12 – 21 maps for future climate conditions); and (3) unsuitable, when < 30% of the maps (up to three maps under current climate and 10 maps under future climates conditions) predicted habitat suitability. We further grouped the habitat suitability maps into two classes by assigning “1” to the pixels that were classified as suitable with high certainty and “0” to the rest to represent suitable and unsuitable habitats, respectively. We overlaid these maps to detect spatiotemporal changes in habitat suitability and quantify the impact of climate change.

### 2.4 Model evaluation

We evaluated the accuracy of each model run by using the receiver operating characteristic curve (ROC), a threshold-independent measure of a model’s ability to discriminate between the pseudo-absence and the presence data [54]. This is a standard method to evaluate the accuracy of predictive distribution models [55] AUC values vary from 0 (random discrimination) to 1 (perfect discrimination) [56]. An AUC value of 0.5 or smaller indicates that the model has no predictive power, whereas perfect discrimination between suitable and unsuitable cells will give an AUC value approaching 1.0 [41].

### 2.5 Historical changes in gelada elevation range

To assess whether the elevation range of geladas already changed in historical times, we compared elevations of historical gelada sightings from the periods before 1900 and before 2000 with the elevations of current sightings (after 2000). We extracted the corresponding elevation range as the difference between maximum and minimum elevation within a radius of four 90-m grid cells around the recorded localities (49 neighbour cells), elevation, and slope standard deviations which are the standard deviation of elevations and slope among these neighbouring grid cells, respectively (ArcGIS version 10.7). We additionally computed slope maximum – the maximum slope among these neighbouring cells and slope range – the difference between maximum and minimum slope values. We further extracted the value of elevation from current and future (2050 and 2070) modelled suitable habitats and compared the change in elevation with the historical data (Fig. S3).

## 3 RESULTS

### 3.1 Variables that predict the distribution of suitable gelada habitat under climate change

Under all settings, mean temperature of the wettest quarter (Bio8), vegetation, slope standard deviation, and precipitation of the wettest month (Bio13) explained most in predicting gelada occurrence (Table 1).

### 3.2 Habitat suitability modeling

All model versions had high predictive performance on both training and test data, with AUC values ≥ 0.87 (Table 2). Models in which a bias file was used for generating pseudo-absence points had relatively lower AUC values (0.88 for the simple model and 0.87 for the complex model), whereas generating pseudo-absence points within the elevation range of gelada, resulted in relatively higher AUC values (0.95 for complex and 0.95 for simple model). No remarkable differences were observed between the AUC values computed on training and test data in the predicted models. The AUC standard deviations of our results demonstrate that there was nearly zero variability or consistency (Std between 0.01 and 0.03), indicating that our data set was accurate enough to make predictions about the suitability of geladas (Table 2). Overall, our prediction was consistent across the model complexity levels, runs, and datasets.

**Table 2.**
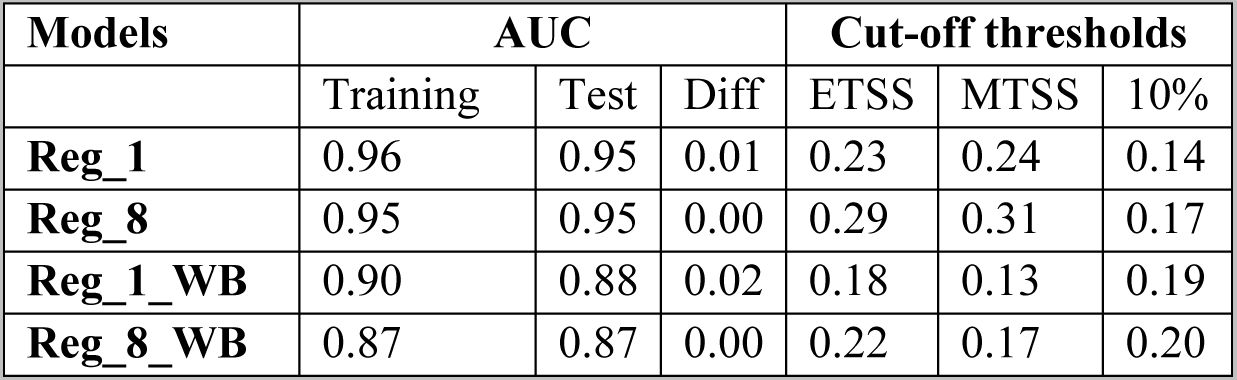
Average training and testing AUC values for the four MaxEnt model versions and their average cut-off of threshold values. Reg_1: complex MaxEnt model run with pseudo-absence points generated within the elevation range of geladas; Reg_8: simple MaxEnt model run with pseudo-absence points generated within the elevation range of geladas; Reg_1_WB: complex MaxEnt model run with pseudo-absence points generated using bias file; Reg_8_WB: simple MaxEnt model run with pseudo-absence points generated using bias file. Diff: the differences between the training and test AUC values.

Under the current climate, the models predicted an area of 90,891 km^2^ to be suitable for geladas (Fig. 3; Table 3). As expected, the suitable habitat mainly concentrates in the northern and central highlands of Ethiopia, where the density of occurrence points is also the highest (Fig. 1). Under future climates conditions, the area predicted to be suitable with high certainty declined to 55,829 km^2^ by 2050 and to 43,576 km^2^ by 2070 (Fig. 3, Table 3, Fig. S4 and S5), a reduction of 39% and 58% by 2050 and 2070, respectively.

**Fig. 3.**
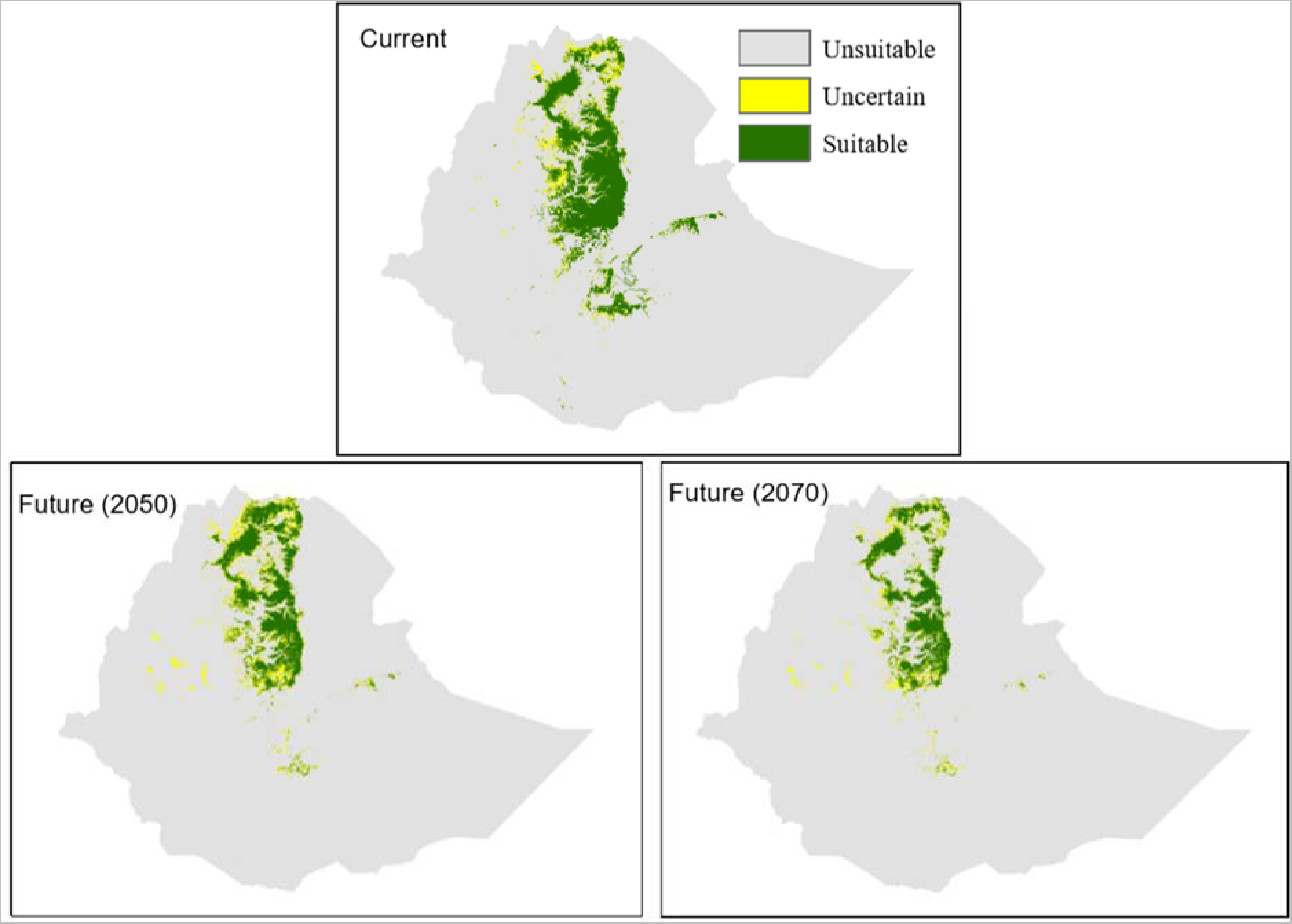
Distribution and extent of suitable gelada habitat produced from 12 binary maps for current climate (current; 2 techniques to generate pseudo-absence points x 2 model complexity levels x 3 threshold values; see also Fig. S2), and from 36 binary maps for future scenarios (future 2050 and 2070); two techniques to generate pseudo-absence points x 2 model complexity levels x 3 threshold values x 3 emission scenarios). When grid cells in 30% or less of the binary maps (3 maps for current and 10 maps for future climates) predict suitability, we considered them unsuitable. When grid cells of >30% - 60% maps (4 – 7 maps for the current climate and 11 – 21 maps for the future climate conditions) predicted suitability, we considered them uncertain in terms of suitability. When grid cells from >60% binary maps (> 7 maps for the current and > 21 maps for the future) predict suitability, we considered them as suitable.

**Table 3.** Loss and gain of suitable habitat for geladas under future climate conditions (2050 and 2070).

| scenarios | current extent<br>km<sup>2</sup> | remain suitable<br>km<sup>2</sup> | loss<br>km<sup>2</sup> | loss<br>% | gain<br>km<sup>2</sup> | gain<br>% | future extent<br>km<sup>2</sup> | change<br>% |
| --- | --- | --- | --- | --- | --- | --- | --- | --- |
| 2050 | 90,891 | 50,362 | 40,529 | 44.6 | 5467 | 6.0 | 55,829 | -38.6 |
| 2070 | 90,891 | 42,696 | 48,195 | 53.0 | 881 | 0.9 | 43,576 | -52.1 |

Under both, current and future climate conditions, the majority of the highly suitable habitat is predicted for the central and northern Ethiopian Highlands (Fig. 3 and 4). The model projections also show some suitable habitat in northern Tigray. In the southern and eastern Ethiopian Highlands, only a few areas with suitable habitat are predicted. In particular, for the Bale, Arsi, and Ahmar Mountains south of the Rift Valley, and for some areas in the central highlands, the models show a loss of habitat. In addition, in these areas, the models indicate not only loss of habitat, but also fragmentation. However, the models also predict a gain in suitable habitat for 2050 and 2070 in northern Ethiopia, specifically in eastern Tigray (Fig. 4 and S5).

**Fig. 4.**
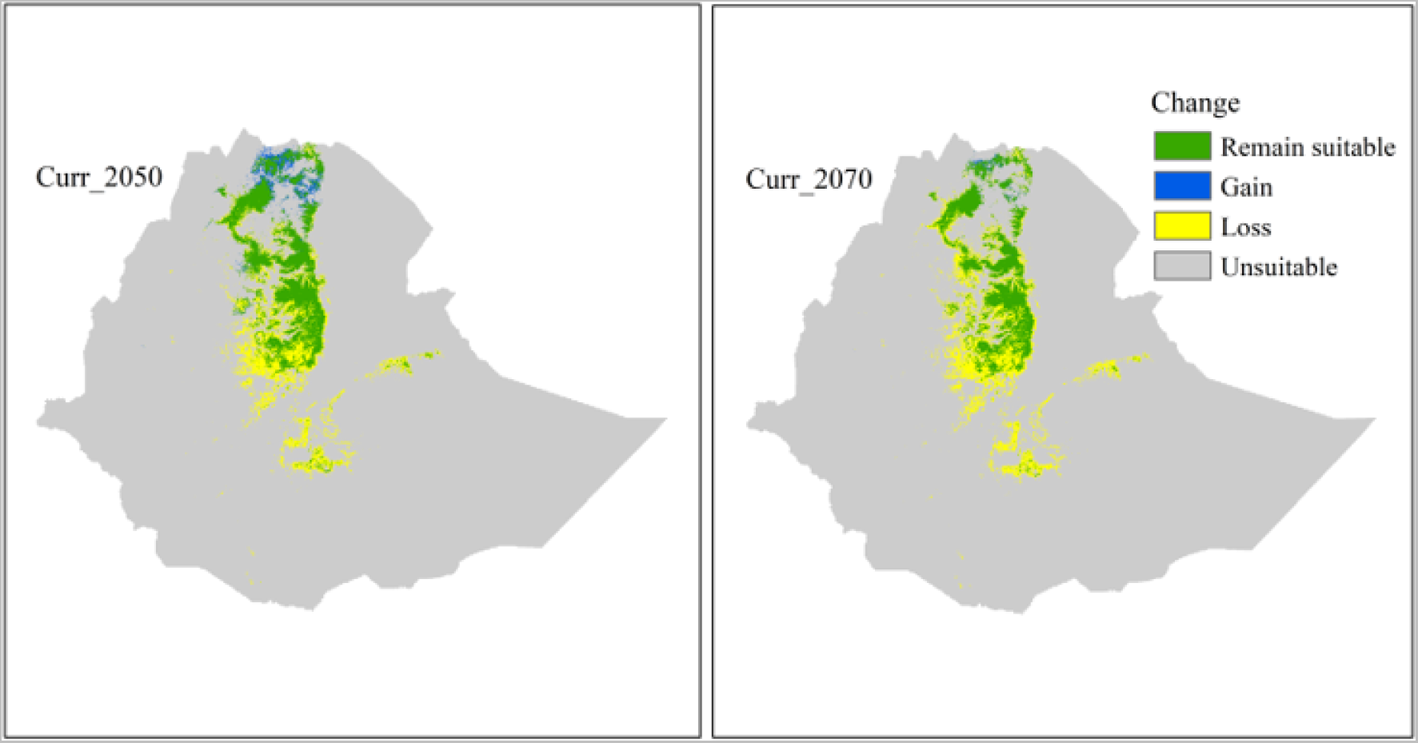
Predicted change in habitat suitablities of geladas by 2050 (Curr_2050) and by 2070 (curr_2070). Green, pixels that are predicted to be suitable under both current and future climates; blue, pixels that are not currently predicted to be suitable but forecasted to be suitable in the future; yellow; currently suitable but not in the future; and grey, unsuitable both under current and future climates.

Elevation and slope of the occurrence points increased over time (1800s to 2000s) (Fig. 5), but elevation did not increase in our projections for 2050 and 2070, compared to the current scenario. The predicted suitable habitat had an average elevation of 2749 m, 2685 m, and 2809 m for the current and future climates, respectively. Similarly, slopes became steeper and the topography more complex over time (Fig. 5).

**Fig. 5.**
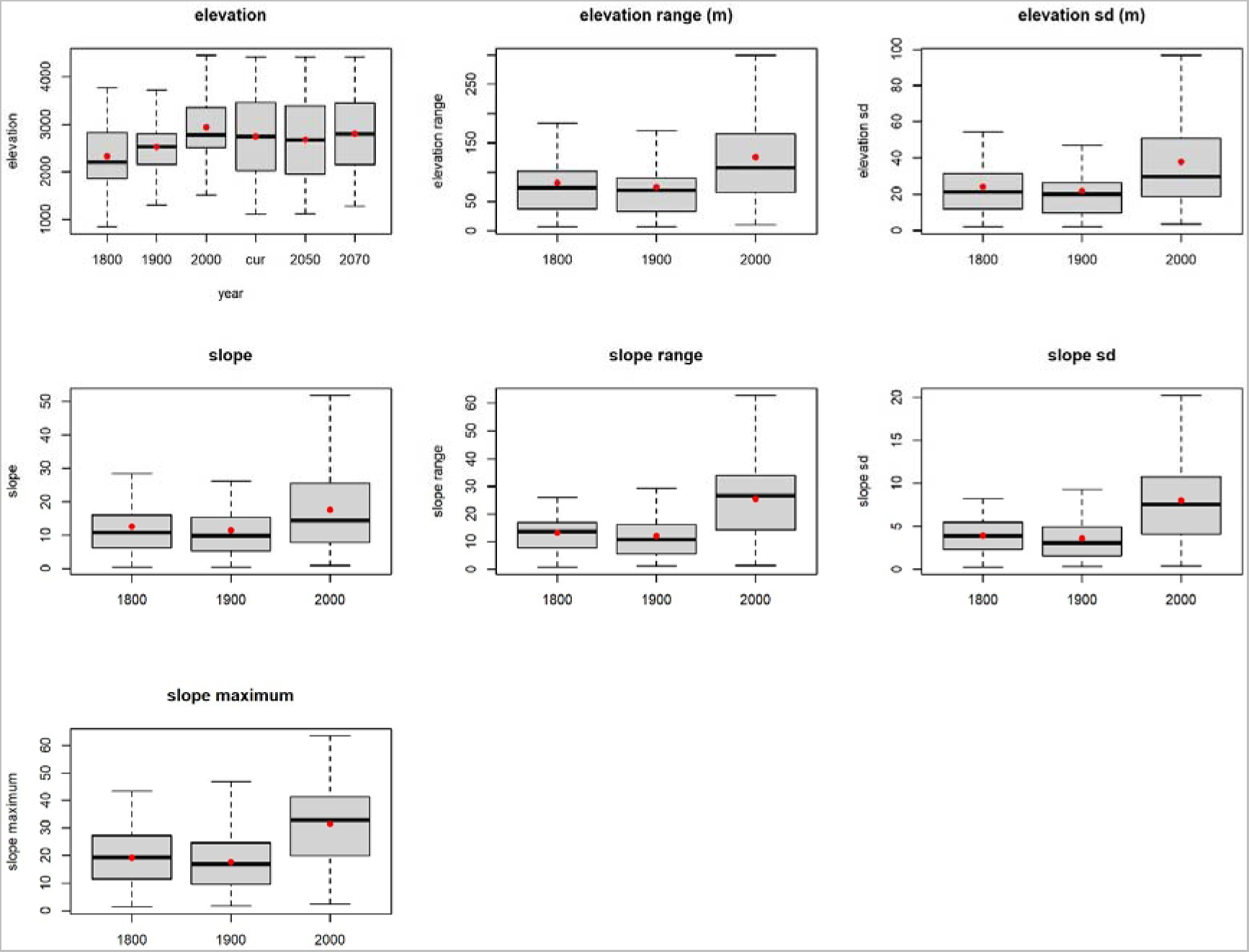
Changes in topographic features (elevation and slope) of gelada occurrence localities during the last 200 years. Box plots depict medians (horizontal lines), quartiles (box), ranges (whiskers), and means (red dots). The temporal variation in elevation of gelada occurrence locations is based on reported sighting from the 1800s, 1900s, and 2000s. For elevation change, we also added elevation of suitable habitat under the current climate (cur), and future climate conditions (2050 and 2070).

The number of pixels per 1 km x 1 km grid cell of suitable habitat varied with elevation (Fig. S3) and the highest number, for all scenarios, was found between 2000 and 3000 m asl. However, the absolute number of pixels under the current (2000) condition was greater than under future scenarios (Fig. S3).

## 4 DISCUSSION

We modelled the distribution of the currently suitable habitat of a high-altitude primate, the gelada, and projected the distribution to future climate scenarios. Our MaxEnt modelling showed high predictive performance for the current distribution and suggests a significant reduction of suitable habitats for geladas under future climates (by 2050 and 2070).

### Modelling

In response to the need for species conservation and management planning in times of climate change, many species distribution modelling approaches have been developed [28, 57]. Though ensemble or model averaging has greater predictive capacity than individual modelling approaches, MaxEnt is commonly used to infer species distributions and environmental tolerances from occurrence data, particularly when optimized well. In this study, we applied different complexity levels, datasets, and cut-off threshold values to tune the predictive performance of the MaxEnt predictions to use the averaged result of our prediction for designing conservation plans for gelada in the Ethiopian highlands.

Although all models had a high performance (AUC > 0.87), the model in which bias files were used to generate pseudo-absence points had a relatively lower AUC, (test AUC = 0.87 for the simple model and 0.88 for the complex model) than the model produced by using pseudo-absence points generated within the elevation range of gelada (0.95 test AUC for both simple and complex models). This may be due to the restriction of randomly generated points in proximity to the presence points when bias file is used. The predicted suitable areas were also lower when bias files were used [48]. Nevertheless, the differences among the modelling approaches were not remarkable. In general, we found consistent results across different model complexities and runs, little difference between test and training AUCs, and similar patterns of prediction among different cut-off thresholds, which indicates the reliability of our approach and the robustness of the models we used. The consistent results across different model complexities and runs indicate acceptable data quality for predictions and the ability of the MaxEnt models to identify the Ethiopian highlands as providing suitable habitat for geladas.

Our modelling shows that the mean temperature of the wettest quarter (Bio 8) was the most influential predictor variable for the distribution of suitable habitat. In a previous study on the consequences of climate change on gelada distribution, Dunbar [26] predicted that geladas will be forced to live only on a few isolated mountain summits if the temperature would increase by 7°C. However, the Intergovernmental Panel on Climate Change [58] estimates that anthropogenically driven climate warming in the 21st century is likely to exceed 1.5°C relative to the 1850–1900 period in all scenarios and exceeds 2.0°C in many scenarios. Though a 7°C temperature increase may not happen within the next 100 years, the result is concerning.

We also found that the distribution of geladas is influenced by annual precipitation and the precipitation of the wettest month. Annual precipitation is associated with food availability and habitat quality [26], and it can affect space useand distribution directly or indirectly by its impact on population dynamics. Results of a recent study on the demography of geladas in the Simien Mountains from 2008 to 2019 suggest that these primates are less resilient to climate variability than previously thought [59].

Although slope had the least average contribution to our models, interestingly slope standard deviation was one of the three most important predictor variables (Table 2). Geladas use the Afro-alpine grasslands on flat plateaus for foraging and steep cliffs as sleeping sites and as refuges in case of predation [16, 60–65]. Slope standard deviation is a good proxy to landscape complexity and thus can capture these topographic niche requirements of gelada as well as other animals with similar adaptations. Thus we recommend the use of slope standard deviation as input especially when distribution models are used to map habitat suitabilities of high-altitude animals.

### Future projection of habitat distribution for *T*. *gelada*

Our averaged model prediction for *T. gelada* shows that the current predicted suitable area covers 90,891 km^2^, while an additional 25,621 km^2^ of potential habitat is considered suitable with uncertainty. Geographically, highly suitable areas were more concentrated in the central highlands, in northern Showa, Wollo, and South Gondar, and the Debre-Libanos area.

Compared to the size of the current suitable habitat, our projections suggested a massive loss of suitable habitats under future climates. Also, small new areas were forecasted to become suitable under climate change in the northern and central parts of Ethiopia, overall the suitable habitat is predicted to decrease by 36% (2050) and 52% (2070), respectively. The most dramatic decline of suitable habitat, however, was projected for the population south of the Rift Valley. Here the size of suitable habitat is already small due to extreme anthropogenic pressure caused by expansions of agriculture, overgrazing, and human-wildlife conflicts as a consequence [22, 23, 66].

### Anthropogenic pressure

Given the strong anthropogenic pressures on gelada habitat overall in Ethiopia, the elevation shift of occurrence points in historical times can most likely be more attributed to agricultural expansion than to the impact of climate change. These pressures at the lower elevation most likely have pushed geladas already to a higher elevation where their climatic resilience might be close to its limit [59]. If geladas are currently living in marginal habitats they might represent a refugee species, which undermines the topographic and climatic tolerances our models predicted. Thus we recommend protecting the current habitats of geladas even when they are forecasted to become climatically unsuitable in the future, in particular for the population south of the Rift Valley. We also recommend conservation efforts even in areas where our models predicted suitable habitats with uncertainty.

## Conclusion

Our species distribution modelling demonstrates that the current suitable habitat of geladas is vulnerable to climate change. Geladas will lose large parts of their current suitable habitat in the Ethiopian highlands. Even though species range shift was not evident in our models, significant elevational changes appeared between current and historical occurrence points, which potentially are associated with anthropogenic pressures at lower elevations. The findings of our study can be used to revisit or align the boundaries of existing protected areas with the future predicted habitats that encompass climate refugia for this high-altitude species. In particular, the population south of the Rift Valley will be severely affected. This is all the more dramatic because no protected areas exist for this (sub)species, thus there is an urgent need to create a protected area for this population.

## Acknowledgments

Addis Ababa University provided facilities to carryout this research. Jens-Christian Svenning was supported by the VILLUM Investigator project “Biodiversity Dynamics in a Changing World”, funded by VILLUM FONDEN (grant 16549), the Centre for Ecological Dynamics in a Novel Biosphere (ECONOVO), funded by the Danish National Research Foundation (grant DNRF173), and the Independent Research Fund Denmark | Natural Sciences project MegaComplexity (grant 0135-00225B).

## Author contributions

ASA and DCG.: data collection and organization, formal analysis, writing (the original draft). CAK: data collection and methodology, organizing, formal analysis, and writing. AA: reviewing, and editing the manuscript. JCS contributes to the comments, editorial and writes a manuscript. AB. edited and reviewed the manuscript, proofread it for the final submission, and is the senior author. DZ. is co-initiator of the study, developed the research question, made significant contributions to, data collection, analysis, and write-up. All authors contributed to the writing of the manuscript. This is original research that has not previously been published and is not under consideration for publication elsewhere. This paper has been approved by all authors.

## Data availability statement

The attached supplementary file contains all of the data.

## Funding

There are no any funding resources for this study.

## Declarations

### Ethics approval and consent to participate

Not applicable

### Consent for publication

Not applicable

### Ethical Guidelines

Not applicable

### Conflict of interest

The authors declare that there is no conflict of interest.

## Supplementary material

**Table S1.** Occurrence locations of gelada (*Theropithecus gelada*)

| # | Region | Taxon | Latitude | Longitude |
| --- | --- | --- | --- | --- |
| 1 | north | *T. g. gelada* | 14.33333 | 39.48333 |
| 2 | north | *T. g. gelada* | 14.16667 | 39.05000 |
| 3 | north | *T. g. gelada* | 14.13333 | 38.71667 |
| 4 | north | *T. g. gelada* | 14.11667 | 39.43333 |
| 5 | north | *T. g. gelada* | 13.87000 | 39.74000 |
| 6 | north | *T. g. gelada* | 13.75000 | 38.53333 |
| 7 | north | *T. g. gelada* | 13.75000 | 38.08333 |
| 8 | north | *T. g. gelada* | 13.71667 | 38.38333 |
| 9 | north | *T. g. gelada* | 13.66667 | 38.41667 |
| 10 | north | *T. g. gelada* | 13.50000 | 38.50000 |
| 11 | north | *T. g. gelada* | 13.49278 | 38.44361 |
| 12 | north | *T. g. gelada* | 13.40000 | 38.20000 |
| 13 | north | *T. g. gelada* | 13.37359 | 38.29285 |
| 14 | north | *T. g. gelada* | 13.35755 | 38.28735 |
| 15 | north | *T. g. gelada* | 13.34038 | 38.09607 |
| 16 | north | *T. g. gelada* | 13.33282 | 38.42881 |
| 17 | north | *T. g. gelada* | 13.30645 | 38.26414 |
| 18 | north | *T. g. gelada* | 13.30402 | 38.42657 |
| 19 | north | *T. g. gelada* | 13.30163 | 38.29574 |
| 20 | north | *T. g. gelada* | 13.30000 | 39.41667 |
| 21 | north | *T. g. gelada* | 13.28391 | 38.12718 |
| 22 | north | *T. g. gelada* | 13.27725 | 38.08100 |
| 23 | north | *T. g. gelada* | 13.27675 | 38.34958 |
| 24 | north | *T. g. gelada* | 13.27604 | 38.09801 |
| 25 | north | *T. g. gelada* | 13.27122 | 38.03430 |
| 26 | north | *T. g. gelada* | 13.27090 | 38.10860 |
| 27 | north | *T. g. gelada* | 13.26603 | 38.14797 |
| 28 | north | *T. g. gelada* | 13.26601 | 38.07775 |
| 29 | north | *T. g. gelada* | 13.26223 | 38.19221 |
| 30 | north | *T. g. gelada* | 13.25511 | 38.20663 |
| 31 | north | *T. g. gelada* | 13.25195 | 38.34669 |
| 32 | north | *T. g. gelada* | 13.25140 | 38.37314 |
| 33 | north | *T. g. gelada* | 13.25000 | 38.25000 |
| 34 | north | *T. g. gelada* | 13.25000 | 38.23333 |
| 35 | north | *T. g. gelada* | 13.25000 | 38.18333 |
| 36 | north | *T. g. gelada* | 13.25000 | 38.15000 |
| 37 | north | *T. g. gelada* | 13.25000 | 38.08333 |
| 38 | north | *T. g. gelada* | 13.25000 | 38.03333 |
| 39 | north | *T. g. gelada* | 13.25000 | 38.00000 |
| 40 | north | *T. g. gelada* | 13.24639 | 37.89528 |
| 41 | north | *T. g. gelada* | 13.24545 | 38.16689 |
| 42 | north | *T. g. gelada* | 13.24078 | 38.36204 |
| 43 | north | *T. g. gelada* | 13.23533 | 38.02050 |
| 44 | north | *T. g. gelada* | 13.23333 | 38.48333 |
| 45 | north | *T. g. gelada* | 13.23333 | 38.41667 |
| 46 | north | *T. g. gelada* | 13.23145 | 38.03955 |
| 47 | north | *T. g. gelada* | 13.23086 | 37.99430 |
| 48 | north | *T. g. gelada* | 13.23063 | 38.06791 |
| 49 | north | *T. g. gelada* | 13.23021 | 38.08360 |
| 50 | north | *T. g. gelada* | 13.22658 | 38.26053 |
| 51 | north | *T. g. gelada* | 13.22439 | 38.02448 |
| 52 | north | *T. g. gelada* | 13.21667 | 38.15000 |
| 53 | north | *T. g. gelada* | 13.21287 | 38.30383 |
| 54 | north | *T. g. gelada* | 13.20899 | 37.99584 |
| 55 | north | *T. g. gelada* | 13.20833 | 38.11667 |
| 56 | north | *T. g. gelada* | 13.20569 | 37.98582 |
| 57 | north | *T. g. gelada* | 13.20300 | 37.88800 |
| 58 | north | *T. g. gelada* | 13.20050 | 38.08333 |
| 59 | north | *T. g. gelada* | 13.19434 | 38.18943 |
| 60 | north | *T. g. gelada* | 13.16667 | 38.00000 |
| 61 | north | *T. g. gelada* | 13.16550 | 38.07260 |
| 62 | north | *T. g. gelada* | 13.15323 | 37.84451 |
| 63 | north | *T. g. gelada* | 13.14319 | 37.84423 |
| 64 | north | *T. g. gelada* | 13.13190 | 37.93228 |
| 65 | north | *T. g. gelada* | 13.12900 | 37.94341 |
| 66 | north | *T. g. gelada* | 13.12636 | 37.83481 |
| 67 | north | *T. g. gelada* | 13.12138 | 37.93323 |
| 68 | north | *T. g. gelada* | 13.11696 | 38.46255 |
| 69 | north | *T. g. gelada* | 13.08200 | 38.43640 |
| 70 | north | *T. g. gelada* | 13.08191 | 38.50800 |
| 71 | north | *T. g. gelada* | 13.06667 | 38.75000 |
| 72 | north | *T. g. gelada* | 13.02274 | 38.10593 |
| 73 | north | *T. g. gelada* | 13.00349 | 38.10598 |
| 74 | north | *T. g. gelada* | 12.98333 | 37.75000 |
| 75 | north | *T. g. gelada* | 12.97711 | 38.35778 |
| 76 | north | *T. g. gelada* | 12.97380 | 38.10574 |
| 77 | north | *T. g. gelada* | 12.83333 | 37.75000 |
| 78 | north | *T. g. gelada* | 12.77207 | 37.55751 |
| 79 | north | *T. g. gelada* | 12.76903 | 37.60704 |
| 80 | north | *T. g. gelada* | 12.70277 | 37.59908 |
| 81 | north | *T. g. gelada* | 12.61667 | 37.45000 |
| 82 | north | *T. g. gelada* | 12.50000 | 37.50000 |
| 83 | central | *T. g. obscurus* | 12.32977 | 38.89366 |
| 84 | central | *T. g. obscurus* | 12.20000 | 37.98333 |
| 85 | central | *T. g. obscurus* | 12.15469 | 39.17117 |
| 86 | central | *T. g. obscurus* | 12.15434 | 39.17168 |
| 87 | central | *T. g. obscurus* | 12.14825 | 39.18215 |
| 88 | central | *T. g. obscurus* | 12.14565 | 39.18215 |
| 89 | central | *T. g. obscurus* | 12.14531 | 39.18227 |
| 90 | central | *T. g. obscurus* | 12.13892 | 39.18552 |
| 91 | central | *T. g. obscurus* | 12.13159 | 39.20464 |
| 92 | central | *T. g. obscurus* | 12.11759 | 39.18675 |
| 93 | central | *T. g. obscurus* | 12.11667 | 39.46667 |
| 94 | central | *T. g. obscurus* | 12.09639 | 39.37812 |
| 95 | central | *T. g. obscurus* | 12.04750 | 39.09624 |
| 96 | central | *T. g. obscurus* | 12.03132 | 38.89378 |
| 97 | central | *T. g. obscurus* | 12.02394 | 39.39683 |
| 98 | central | *T. g. obscurus* | 12.01667 | 39.05000 |
| 99 | central | *T. g. obscurus* | 12.00000 | 39.00000 |
| 100 | central | *T. g. obscurus* | 11.86372 | 39.19617 |
| 101 | central | *T. g. obscurus* | 11.83333 | 38.08333 |
| 102 | central | *T. g. obscurus* | 11.81667 | 38.65786 |
| 103 | central | *T. g. obscurus* | 11.81257 | 38.67056 |
| 104 | central | *T. g. obscurus* | 11.81100 | 38.69572 |
| 105 | central | *T. g. obscurus* | 11.80884 | 38.68606 |
| 106 | central | *T. g. obscurus* | 11.73286 | 38.89390 |
| 107 | central | *T. g. obscurus* | 11.73186 | 38.18903 |
| 108 | central | *T. g. obscurus* | 11.70341 | 38.23743 |
| 109 | central | *T. g. obscurus* | 11.69748 | 39.25106 |
| 110 | central | *T. g. obscurus* | 11.60368 | 38.92768 |
| 111 | central | *T. g. obscurus* | 11.59543 | 38.94091 |
| 112 | central | *T. g. obscurus* | 11.55421 | 39.22692 |
| 113 | central | *T. g. obscurus* | 11.55181 | 39.10517 |
| 114 | central | *T. g. obscurus* | 11.53333 | 39.21667 |
| 115 | central | *T. g. obscurus* | 11.51261 | 39.01186 |
| 116 | central | *T. g. obscurus* | 11.49615 | 38.93973 |
| 117 | central | *T. g. obscurus* | 11.48385 | 38.79960 |
| 118 | central | *T. g. obscurus* | 11.47721 | 38.97112 |
| 119 | central | *T. g. obscurus* | 11.46081 | 38.82572 |
| 120 | central | *T. g. obscurus* | 11.45757 | 38.84944 |
| 121 | central | *T. g. obscurus* | 11.45091 | 39.24459 |
| 122 | central | *T. g. obscurus* | 11.44982 | 38.99535 |
| 123 | central | *T. g. obscurus* | 11.44982 | 38.94951 |
| 124 | central | *T. g. obscurus* | 11.43439 | 38.89401 |
| 125 | central | *T. g. obscurus* | 11.43333 | 39.31667 |
| 126 | central | *T. g. obscurus* | 11.38629 | 39.24910 |
| 127 | central | *T. g. obscurus* | 11.36742 | 39.24329 |
| 128 | central | *T. g. obscurus* | 11.25000 | 39.58333 |
| 129 | central | *T. g. obscurus* | 11.23220 | 39.21514 |
| 130 | central | *T. g. obscurus* | 11.16983 | 39.24077 |
| 131 | central | *T. g. obscurus* | 11.11667 | 39.71667 |
| 132 | central | *T. g. obscurus* | 11.11210 | 39.14120 |
| 133 | central | *T. g. obscurus* | 11.07247 | 39.27562 |
| 134 | central | *T. g. obscurus* | 11.06100 | 39.66312 |
| 135 | central | *T. g. obscurus* | 11.06007 | 39.66022 |
| 136 | central | *T. g. obscurus* | 11.05917 | 39.66114 |
| 137 | central | *T. g. obscurus* | 11.05809 | 39.66176 |
| 138 | central | *T. g. obscurus* | 10.95747 | 38.68090 |
| 139 | central | *T. g. obscurus* | 10.95197 | 38.66504 |
| 140 | central | *T. g. obscurus* | 10.91760 | 38.79465 |
| 141 | central | *T. g. obscurus* | 10.89623 | 38.85149 |
| 142 | central | *T. g. obscurus* | 10.89421 | 38.80419 |
| 143 | central | *T. g. obscurus* | 10.89290 | 38.82110 |
| 144 | central | *T. g. obscurus* | 10.88725 | 37.51831 |
| 145 | central | *T. g. obscurus* | 10.86667 | 38.75000 |
| 146 | central | *T. g. obscurus* | 10.85150 | 38.79275 |
| 147 | central | *T. g. obscurus* | 10.85028 | 38.68679 |
| 148 | central | *T. g. obscurus* | 10.74129 | 39.18177 |
| 149 | central | *T. g. obscurus* | 10.73444 | 38.78564 |
| 150 | central | *T. g. obscurus* | 10.73353 | 38.88564 |
| 151 | central | *T. g. obscurus* | 10.72312 | 39.29682 |
| 152 | central | *T. g. obscurus* | 10.69457 | 39.32644 |
| 153 | central | *T. g. obscurus* | 10.69262 | 39.36281 |
| 154 | central | *T. g. obscurus* | 10.68437 | 39.18207 |
| 155 | central | *T. g. obscurus* | 10.67874 | 39.71903 |
| 156 | central | *T. g. obscurus* | 10.67739 | 39.23229 |
| 157 | central | *T. g. obscurus* | 10.67483 | 39.31803 |
| 158 | central | *T. g. obscurus* | 10.66667 | 37.95000 |
| 159 | central | *T. g. obscurus* | 10.65850 | 39.41592 |
| 160 | central | *T. g. obscurus* | 10.65739 | 39.25292 |
| 161 | central | *T. g. obscurus* | 10.65000 | 37.75000 |
| 162 | central | *T. g. obscurus* | 10.63899 | 39.80591 |
| 163 | central | *T. g. obscurus* | 10.63879 | 39.13340 |
| 164 | central | *T. g. obscurus* | 10.63422 | 39.17714 |
| 165 | central | *T. g. obscurus* | 10.61601 | 39.33640 |
| 166 | central | *T. g. obscurus* | 10.61250 | 39.68081 |
| 167 | central | *T. g. obscurus* | 10.58771 | 39.63434 |
| 168 | central | *T. g. obscurus* | 10.58753 | 39.44739 |
| 169 | central | *T. g. obscurus* | 10.56086 | 39.59470 |
| 170 | central | *T. g. obscurus* | 10.55490 | 39.44455 |
| 171 | central | *T. g. obscurus* | 10.55338 | 39.68351 |
| 172 | central | *T. g. obscurus* | 10.54603 | 39.56152 |
| 173 | central | *T. g. obscurus* | 10.50315 | 39.54473 |
| 174 | central | *T. g. obscurus* | 10.50307 | 39.57781 |
| 175 | central | *T. g. obscurus* | 10.47395 | 39.51856 |
| 176 | central | *T. g. obscurus* | 10.45169 | 39.17690 |
| 177 | central | *T. g. obscurus* | 10.42464 | 39.80154 |
| 178 | central | *T. g. obscurus* | 10.41404 | 39.74936 |
| 179 | central | *T. g. obscurus* | 10.41124 | 39.77116 |
| 180 | central | *T. g. obscurus* | 10.40927 | 39.26444 |
| 181 | central | *T. g. obscurus* | 10.39131 | 39.38805 |
| 182 | central | *T. g. obscurus* | 10.39125 | 39.41584 |
| 183 | central | *T. g. obscurus* | 10.36555 | 39.48019 |
| 184 | central | *T. g. obscurus* | 10.35000 | 39.78333 |
| 185 | central | *T. g. obscurus* | 10.33700 | 39.47710 |
| 186 | central | *T. g. obscurus* | 10.32744 | 39.80490 |
| 187 | central | *T. g. obscurus* | 10.32593 | 39.20831 |
| 188 | central | *T. g. obscurus* | 10.31758 | 39.80493 |
| 189 | central | *T. g. obscurus* | 10.31383 | 39.81494 |
| 190 | central | *T. g. obscurus* | 10.30559 | 39.29125 |
| 191 | central | *T. g. obscurus* | 10.30094 | 39.81064 |
| 192 | central | *T. g. obscurus* | 10.30018 | 39.19937 |
| 193 | central | *T. g. obscurus* | 10.29453 | 39.25761 |
| 194 | central | *T. g. obscurus* | 10.29119 | 39.78599 |
| 195 | central | *T. g. obscurus* | 10.26238 | 39.08645 |
| 196 | central | *T. g. obscurus* | 10.25106 | 39.07492 |
| 197 | central | *T. g. obscurus* | 10.25000 | 40.00000 |
| 198 | central | *T. g. obscurus* | 10.23347 | 39.11314 |
| 199 | central | *T. g. obscurus* | 10.23256 | 39.17198 |
| 200 | central | *T. g. obscurus* | 10.22663 | 39.09321 |
| 201 | central | *T. g. obscurus* | 10.22656 | 39.15349 |
| 202 | central | *T. g. obscurus* | 10.22345 | 39.39972 |
| 203 | central | *T. g. obscurus* | 10.21279 | 39.09042 |
| 204 | central | *T. g. obscurus* | 10.19060 | 39.00205 |
| 205 | central | *T. g. obscurus* | 10.09447 | 39.49560 |
| 206 | central | *T. g. obscurus* | 10.08333 | 38.28333 |
| 207 | central | *T. g. obscurus* | 10.06673 | 38.21131 |
| 208 | central | *T. g. obscurus* | 10.06667 | 38.28333 |
| 209 | central | *T. g. obscurus* | 10.06617 | 39.02085 |
| 210 | central | *T. g. obscurus* | 10.06308 | 39.60314 |
| 211 | central | *T. g. obscurus* | 9.93466 | 39.23103 |
| 212 | central | *T. g. obscurus* | 9.92401 | 39.12938 |
| 213 | central | *T. g. obscurus* | 9.92128 | 38.92689 |
| 214 | central | *T. g. obscurus* | 9.91667 | 39.78333 |
| 215 | central | *T. g. obscurus* | 9.90000 | 39.78333 |
| 216 | central | *T. g. obscurus* | 9.84291 | 38.90653 |
| 217 | central | *T. g. obscurus* | 9.83989 | 38.89557 |
| 218 | central | *T. g. obscurus* | 9.83854 | 39.74226 |
| 219 | central | *T. g. obscurus* | 9.83333 | 39.78333 |
| 220 | central | *T. g. obscurus* | 9.82280 | 38.88168 |
| 221 | central | *T. g. obscurus* | 9.82199 | 39.70896 |
| 222 | central | *T. g. obscurus* | 9.82071 | 39.73438 |
| 223 | central | *T. g. obscurus* | 9.81673 | 38.89802 |
| 224 | central | *T. g. obscurus* | 9.81168 | 38.73698 |
| 225 | central | *T. g. obscurus* | 9.80000 | 38.75000 |
| 226 | central | *T. g. obscurus* | 9.79116 | 39.68223 |
| 227 | central | *T. g. obscurus* | 9.78930 | 38.99210 |
| 228 | central | *T. g. obscurus* | 9.78436 | 38.95766 |
| 229 | central | *T. g. obscurus* | 9.77978 | 39.75060 |
| 230 | central | *T. g. obscurus* | 9.75675 | 38.85727 |
| 231 | central | *T. g. obscurus* | 9.75000 | 39.75000 |
| 232 | central | *T. g. obscurus* | 9.74411 | 38.86093 |
| 233 | central | *T. g. obscurus* | 9.73816 | 39.73623 |
| 234 | central | *T. g. obscurus* | 9.73742 | 38.81266 |
| 235 | central | *T. g. obscurus* | 9.73190 | 39.74961 |
| 236 | central | *T. g. obscurus* | 9.72799 | 38.82169 |
| 237 | central | *T. g. obscurus* | 9.71682 | 38.83738 |
| 238 | central | *T. g. obscurus* | 9.71667 | 38.86667 |
| 239 | central | *T. g. obscurus* | 9.71519 | 38.84727 |
| 240 | central | *T. g. obscurus* | 9.70330 | 38.88094 |
| 241 | central | *T. g. obscurus* | 9.70000 | 39.50000 |
| 242 | central | *T. g. obscurus* | 9.70000 | 38.81667 |
| 243 | central | *T. g. obscurus* | 9.67276 | 39.50965 |
| 244 | central | *T. g. obscurus* | 9.66667 | 39.53333 |
| 245 | central | *T. g. obscurus* | 9.66667 | 39.05000 |
| 246 | central | *T. g. obscurus* | 9.66667 | 39.03333 |
| 247 | central | *T. g. obscurus* | 9.65000 | 39.75000 |
| 248 | central | *T. g. obscurus* | 9.63333 | 39.31667 |
| 249 | central | *T. g. obscurus* | 9.62176 | 39.73863 |
| 250 | central | *T. g. obscurus* | 9.58333 | 39.75000 |
| 251 | central | *T. g. obscurus* | 9.57642 | 38.91767 |
| 252 | central | *T. g. obscurus* | 9.51667 | 38.21667 |
| 253 | central | *T. g. obscurus* | 9.50000 | 38.16667 |
| 254 | central | *T. g. obscurus* | 9.49340 | 38.43410 |
| 255 | central | *T. g. obscurus* | 9.48000 | 38.43000 |
| 256 | central | *T. g. obscurus* | 9.46667 | 38.75000 |
| 257 | central | *T. g. obscurus* | 9.45260 | 39.43200 |
| 258 | central | *T. g. obscurus* | 9.44060 | 38.48510 |
| 259 | central | *T. g. obscurus* | 9.43496 | 39.53940 |
| 260 | central | *T. g. obscurus* | 9.43473 | 38.65323 |
| 261 | central | *T. g. obscurus* | 9.43010 | 38.50060 |
| 262 | central | *T. g. obscurus* | 9.41667 | 38.75000 |
| 263 | central | *T. g. obscurus* | 9.30917 | 38.73402 |
| 264 | central | *T. g. obscurus* | 9.30000 | 38.61667 |
| 265 | central | *T. g. obscurus* | 9.25000 | 38.55000 |
| 266 | central | *T. g. obscurus* | 9.16667 | 39.41667 |
| 267 | central | *T. g. obscurus* | 9.13333 | 39.10000 |
| 268 | central | *T. g. obscurus* | 9.13333 | 39.03333 |
| 269 | central | *T. g. obscurus* | 9.11667 | 39.11667 |
| 270 | central | *T. g. obscurus* | 9.08333 | 38.75000 |
| 271 | central | *T. g. obscurus* | 8.91667 | 38.61667 |
| 272 | central | *T. g. obscurus* | 8.85412 | 38.84637 |
| 273 | south | *T. g. ssp. nov* | 7.90306 | 39.27167 |
| 274 | south | *T. g. ssp. nov* | 7.74965 | 39.81547 |
| 275 | south | *T. g. ssp. nov* | 7.73912 | 39.83617 |
| 276 | south | *T. g. ssp. nov* | 7.70422 | 39.81851 |
| 277 | south | *T. g. ssp. nov* | 7.69513 | 39.81240 |
| 278 | south | *T. g. ssp. nov* | 7.69358 | 39.82556 |
| 279 | south | *T. g. ssp. nov* | 7.68000 | 40.18300 |
| 280 | south | *T. g. ssp. nov* | 7.53947 | 39.94557 |
| 281 | south | *T. g. ssp. nov* | 7.52680 | 40.02117 |
| 282 | south | *T. g. ssp. nov* | 7.51617 | 39.96753 |
| 283 | south | *T. g. ssp. nov* | 7.50983 | 39.99065 |
| 284 | south | *T. g. ssp. nov* | 7.50330 | 39.51300 |
| 285 | south | *T. g. ssp. nov* | 7.50055 | 39.99195 |

**FIGURE S1.**
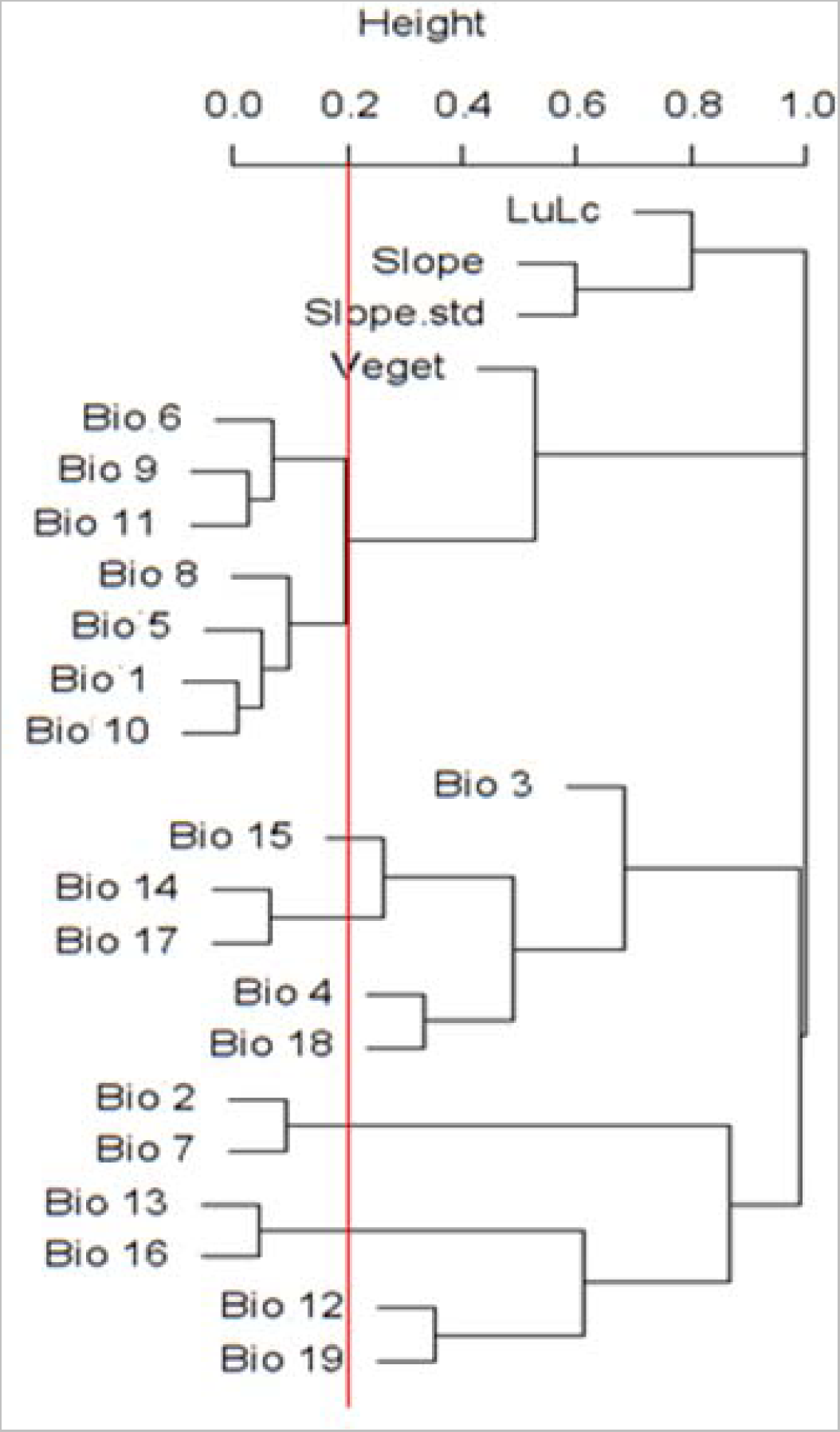
Pairwise Pearson correlation of the predictor variables at locations of training and evaluation datasets. Redline shows a correlation coefficient r = |0.8|. Annual Mean Temperature (Bio1), Mean Diurnal Range (mean of monthly max temp - min temp; Bio2), Isothermality (Bio3), Temperature Seasonality (Bio4), Maximum Temperature of Warmest Month (Bio5), Minimum Temperature of Coldest Month (Bio6), Temperature Annual Range (Bio7), Mean Temperature of Wettest Quarter (Bio8), Annual Precipitation (Bio12), Precipitation of Wettest Month (Bio13), Precipitation of Driest Month (Bio14), Precipitation Seasonality (Coefficient of Variation) (Bio15), Precipitation of Wettest Quarter (Bio16), Precipitation of Driest Quarter (Bio17), Precipitation of Warmest Quarter (Bio18), Precipitation of Coldest Quarter (Bio19), Slope, Slope Standard Deviation (Slope. Std), Land use land cover change (LuLc) and Vegetation.

**Figure S2.**
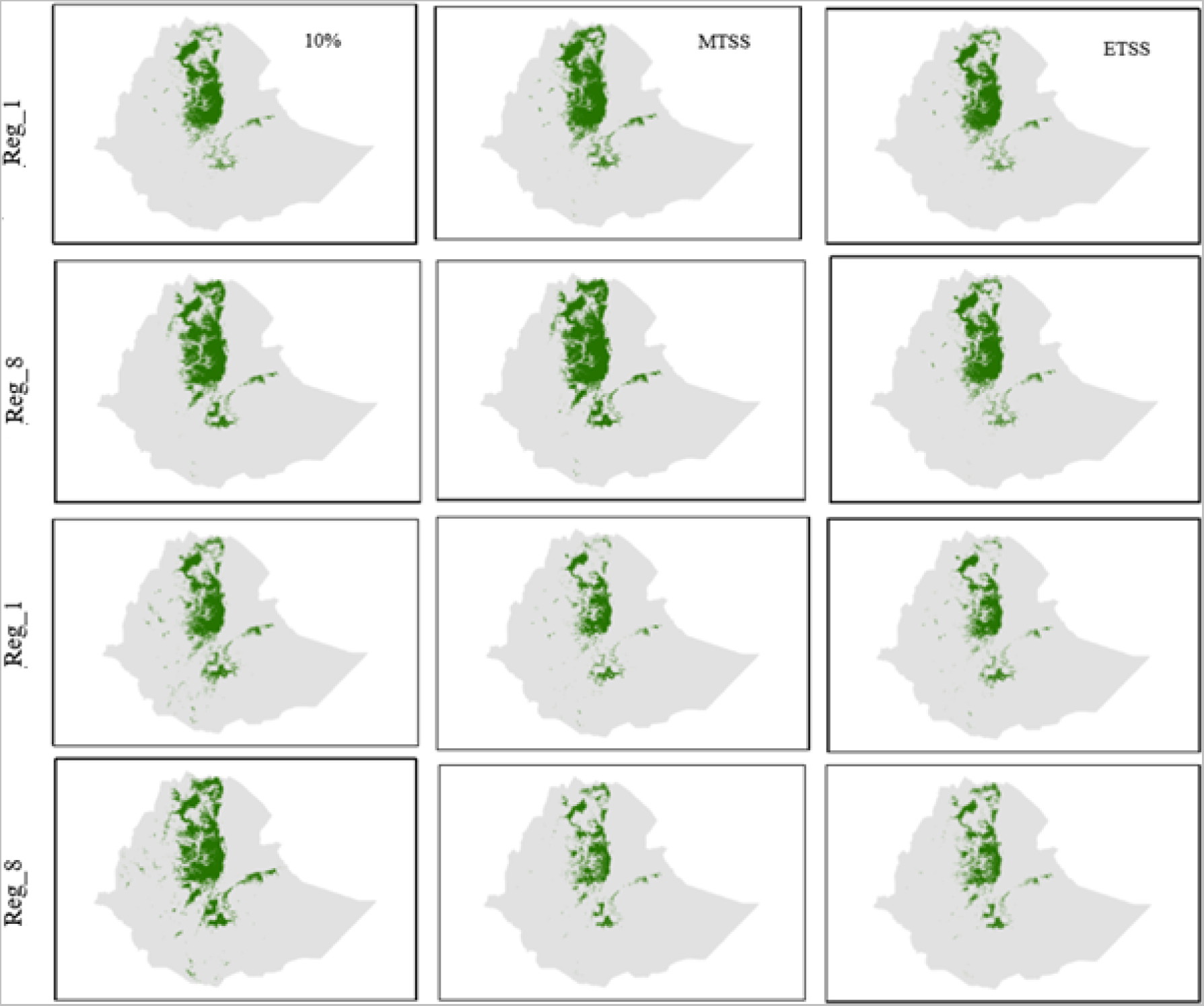
Distribution and extent of suitable habitat of *T*. *gelada* produced using two levels of model complexity (regularization multiplier value = 1 (Reg_1) and 8 (Reg_8)) and three cut-off threshold values: 10% (10 percent omission rate), MTSS (maximum test sensitivity and specificity), and ETSS (equal test sensitivity and specificity). The maps in the upper two rows were produced by generating pseudoabsence points within the elevation ranges of the occurrence points of *T*. *gelada* (2018 – 4219 m asl), while the lower two rows were produced by generating pseudoabsence points using a bias file.

**FIGURE S3.**
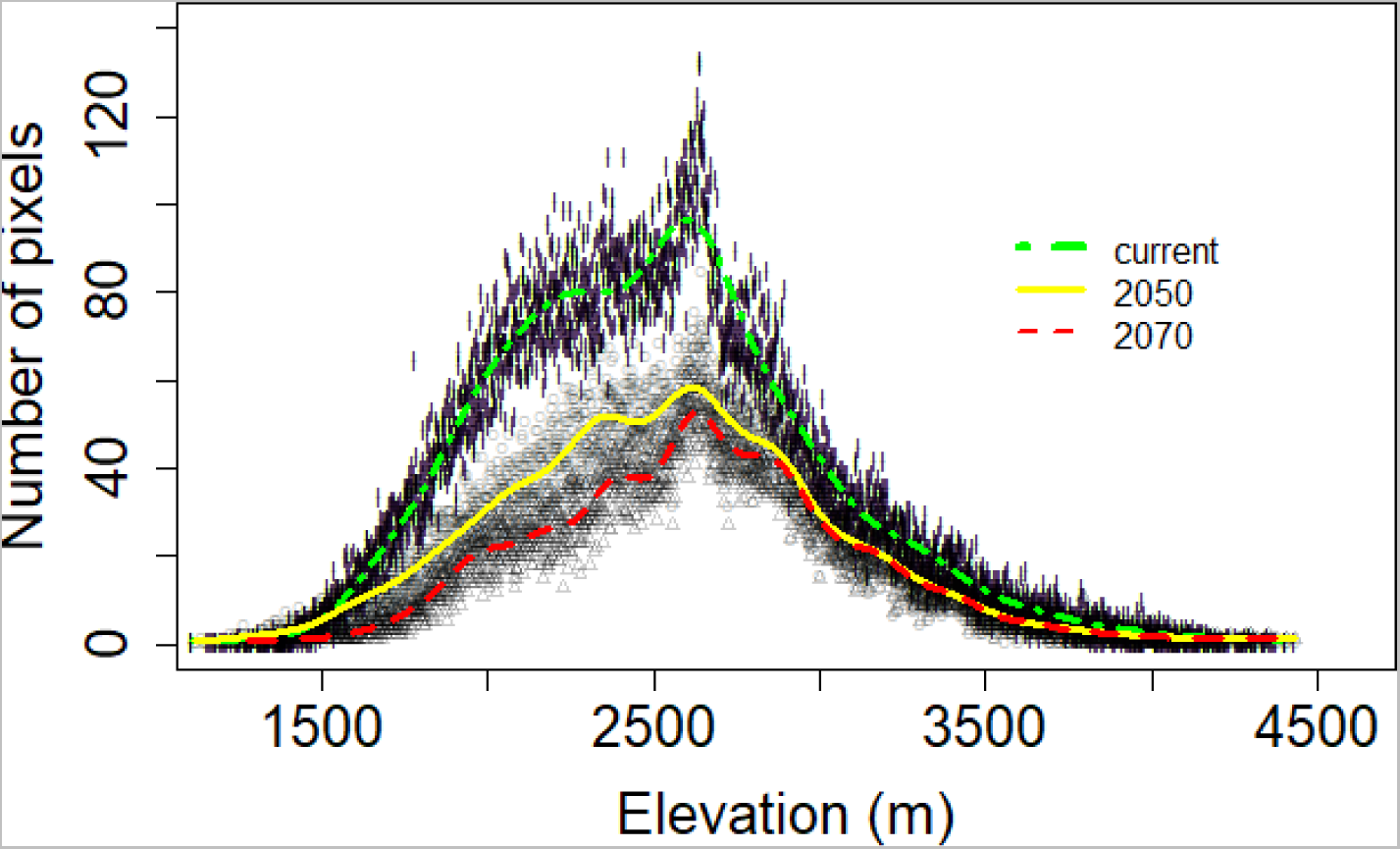
Predicted size of suitable habitats of gelada across an elevation gradient under current and future climates (2050 and 2070). The x-axis represents altitude (m) and the y-axis represents 1 km × 1 km grid cell counts.

**FIGURE S4.**
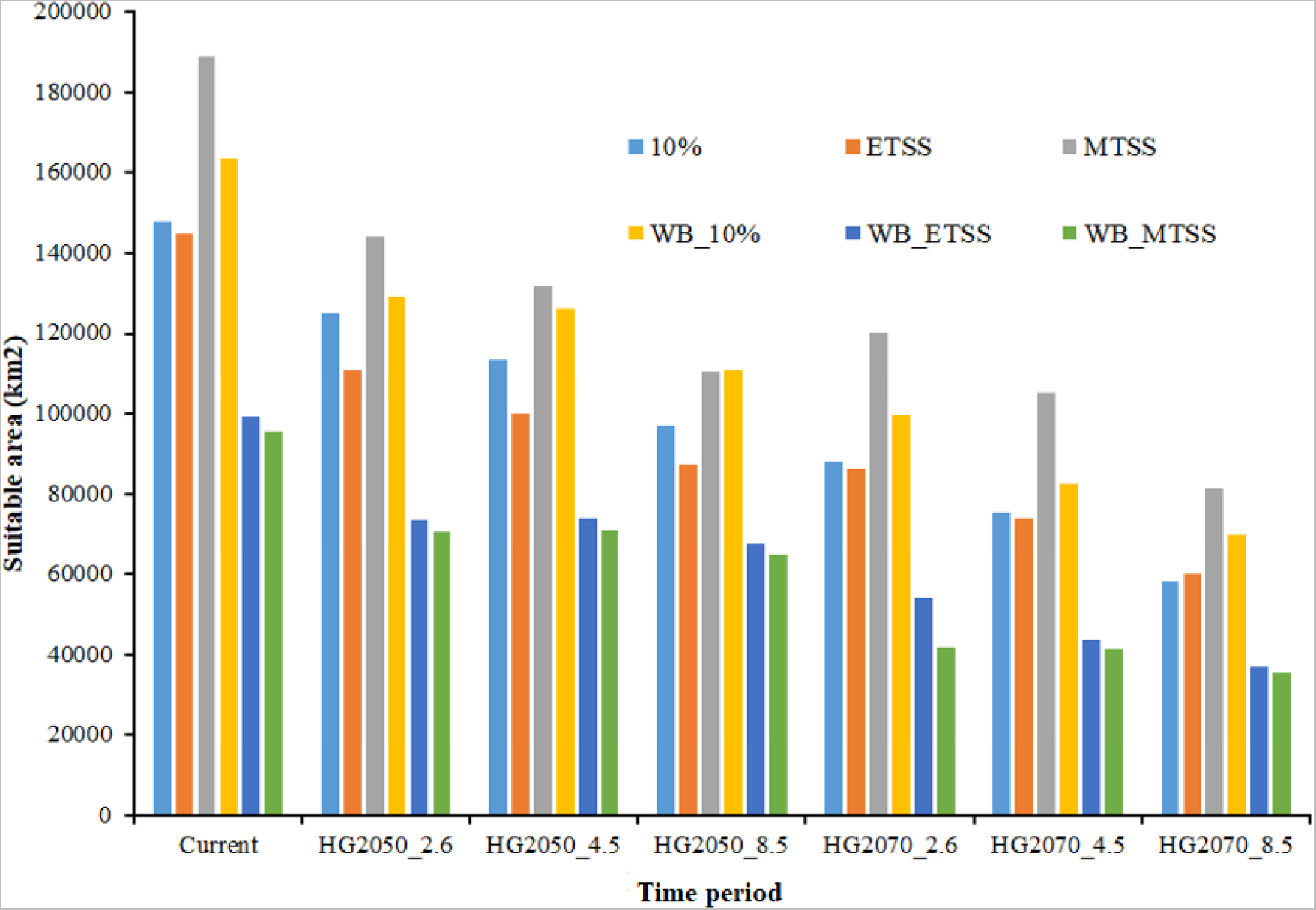
The area of suitable gelada habitats predicted under current and different future emission scenarios (2050 and 2070) produced using two levels of model complexity by setting a regularization multiplier values to 1 (Reg_1) and 8 (Reg_8) and by using three cut-off threshold values without and with bias file (WB): 10% (10 percentile omission rate), MTSS (maximum test sensitivity and specificity), and ETSS (equal test sensitivity and specificity). HG stands for Hadley Centre Global Environment Model version 2 (HadGEM2-ES). We used three emission scenarios (2.6, 4.5 and 8.5).

**FIGURE S5.**
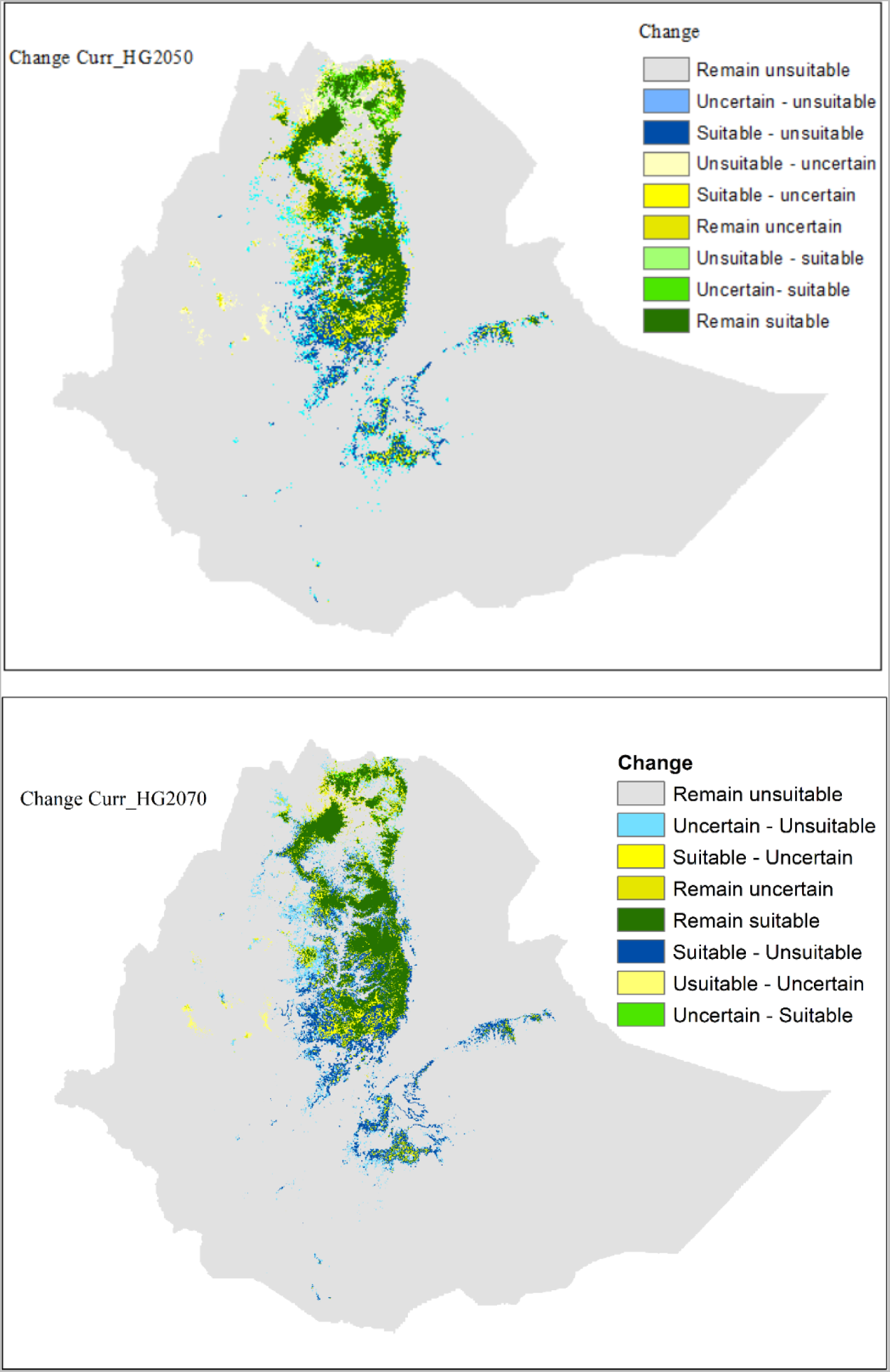
Changes in distribution and extent of suitable gelada habitat. The habitat suitability maps show predicted losses and gains by comparing current and future projections (curr_2050 and curr_2070).

## Notes

### Competing Interest Statement

The authors have declared no competing interest.

